# Rational selection and Characterisation of bile acid (BA) metabolising species of infant origin

**DOI:** 10.1101/2022.06.24.497474

**Authors:** Sarah L. Long, Susan A. Joyce

## Abstract

Bile acids (BAs), biological detergents for nutrient digestion, are important local and systemic signalling molecules to interact with a variety of cell receptors central to influence host responses. While BAs are synthesized in the liver, the range and diversity of bile acids available to interact with these receptors is dictated by the gut microbiota. Bile salt hydrolase (BSH) activity is one such function, it is commonly represented and highly conserved across all major bacterial phyla in the gut. Studies relating to the importance of such modifications in early life are scarce. This study highlights BA metabolism diversity by functionally isolating BA metabolizing strains and by characterizing specific classes of BSH from the formula–fed transitioning gut. Isolates were identified to species levels, *in silico* and *in vitro* characterisation of their BSH genetic content, enzyme activity and substrate specificity. One of these isolates was identified as *Lactobacillus acidophilus*, a species frequently applied as a probiotic whereas three of these four isolates were identified as *Enterococcus avium*. This particular species is not well characterized in the literature and to our knowledge this is the first report of BSH activity and assessment for probiotic potential within this class of microbes. This study indicates that microbial BA altering activity appears functionally reduced, in the formula fed infant gut.

## Introduction

Microbial gut colonization in the first years of life is a critical and influential factor that can influence health in later life (Yatsunenko *et al*., 2012). Numerous infant studies have identified key bacterial successions and implicate these successions as important for gut maturation and homeostasis (Yatsunenko *et al*., 2012). The gut microbiota is relatively unstable at birth but appears to stabilize by three to four years of age unless impacted upon by antibiotic use, injury or state of health (Rutayisire *et al*., 2016, Yassour *et al*., 2016). The infant gut ecosystem appears to follow a sequence of natural succession. The first invaders include facultative anaerobes of the phylum *Actinobacteria, Firmicutes* and *Proteobacteria* characterized by a range of *Enterococcal* and *E. coli* species respectively, these are followed by *Firmicute* colonisation to include *Clostridium* and *Bifidobacterium* species (both obligate anaerobes). Interestingly the degree of colonization and the change in succession appears dependent on mode of birth, gestational period and, in particular, pivotal points of dietary change (Bokulich *et al*., 2016, Yassour *et al*., 2016, Tanaka and Nakayama 2017) and reviewed by (Rutayisire *et al*., 2016).

*Proteobacteria, Bacteroides, Firmicutes (Clostridium)* and Actinobacteria (*Bifidobacteria* species) feature as early infant gut colonizers following normal vaginal birth, the latter is reduced in infants born through C section. This reduction is associated with decreased colonization resistance and increased microbial diversity in early life (Huurre *et al*., 2008). Interestingly, all of these phyla contain bacterial species that are capable of modifying bile acids through bile salt hydrolase activity to remove the amide from liver produced bile acids (deconjugation) (see (Liang *et al*., 2018)). Some members of *Clostridia* IXVa can then dehydroxylate these freed BA to produce secondary BA production (deoxycholic acid (DCA) from free cholic acid (CA), lithocholic acid (LCA) from free chenodeoxycholic acid (CDCA) while others produce ursodeoxycholic acid (UDCA) from CDCA (see review by (Long *et al*., 2017a)). Taken together, the gut microbiota are responsible for the BA diverse palate derived from liver produced BAs that function to emulsify dietary lipids, induce hormones and lipid digesting enzymes, act as differential signalling molecules to alter metabolic processes, influence circadian circuitry as well as immune processes (Joyce *et al*., 2014, Albaugh *et al*., 2019).

*Bifidobacteria* and *Lactobacillus*, both lactic acid bacteria (LAB) species, are particularly potent BA deconjugators (Song *et al*., 2019). They appear earlier, and at higher levels, in breast fed over formula fed infants (Cooke *et al*., 2005, Lewis *et al*., 2017). Formula feeding is associated with elevations in *Enterococcus, Escherichia coli* and *Staphlococcus* species Interestingly, Nagpal *et al*., (2018) isolated a range of breast fed, infant, faecal bacterial isolates including LAB and Enterococcal species that could contribute to SCFA enrichment. Their ability to tolerate bile was assessed, although modifications of bile acids were not addressed (Nagpal *et al*., 2018). Furthermore, bile acids, CA and CDCA, are present breast milk (Forsyth *et al*., 1983). Indeed, UDCA treatment for mothers with biliary cirrhosis could elevate the levels of LCA but not UDCA in human breast milk (Vítek *et al*., 2010), suggesting selective bile acid transport into breast milk. The role of bile acids in milk is suspected to enhance microbial colonization resistance while enhancing milk fat emulsification, although neither are proven. It is also possible that these BA moieties may influence gut-liver signalling, regional cell development and gut homeostasis.

A pre-requisite for probiotic strain selection is the ability to metabolize BA in order to survive the stresses it presents in the GI tract. Indeed, probiotic strains, mainly LAB, have been isolated from normal birthed and breast fed infants and include many different *Lactobacillus* and *Bifidobacteria* strains (Kirtzalidou *et al*., 2011, Muñoz-Quezada *et al*., 2013, Panya *et al*., 2016, Deshpande *et al*., 2017, Lenfestey and Neu 2017, O. Ogunshe 2017). They have been applied to treat a range of paediatric conditions with some success including colic (Sung *et al*., 2018), necrotizing enterocolitis (Manzoni *et al*., 2014) allergenic response modulation (Berni Canani *et al*., 2017) and for promoting general gut health (Radke *et al*., 2017). The body of evidence supporting the role of these microbially altered BAs and their differential effects on cell metabolism through NRs, GCPR and transporters (reviewed in Long *et al*., 2017) points to a central role for microbial BA metabolism in nutrition and a potential role for BAs in gut development and homeostasis. Further insights into the BA modifications by early colonising bacteria, on the basis of diet, en route to a relatively stable microbiome could inform future work in the area of infant gut development.

Here, we functionally isolated, classified and genetically examined a range of BA metabolising gut bacteria from a formula fed 2 year old, representing the transitioning gut. Combining and developing a range of genetic analysis tools specific *Lactobacillus* and *Enterococcus* species, were identified to the strain level. This functional screen revealed representatives of a relatively uncharacterized bacterial species, *Enterococcus avium*. We examined strain collective and gene specific BA modification ability to reveal known as well as novel uncharacterized BSH enzymes for *Lactobacillus* and *E. avium* species respectively. Surprisingly, our data indicate that BA modification was maintained at a low level with restricted substrate ranges associated with individual metabolizing enzymes. We further investigated their potential to be applied as probiotics according to EFSA guidelines. To our knowledge this is the first report and in-depth characterisation of functional BA metabolism based screening and specific BSH activity assessment in the transitioning gut. It is also the first report to examine the probiotic potential and develop a MLST based classification system for *Enterococcus avium*.

## Results

### Functional screening for bile metabolizing bacteria of infant origin

In total, 792 isolates were subjected to bile acid deconjugation assay using agar media containing one of TCA, TDCA and GDCA. Activity was scored as positive where a white precipitate was visible indicating release of hydrophobic bile acids (CA or DCA) (Figure 1). From this assay 70% of isolates (representing 588 colonies) displayed bile metabolising activity however only 7.5% of these isolates (representing 60 colonies) were capable of deconjugating all bile acids tested. These 60 isolates were carried forward for further analysis.

**Figure 1:**
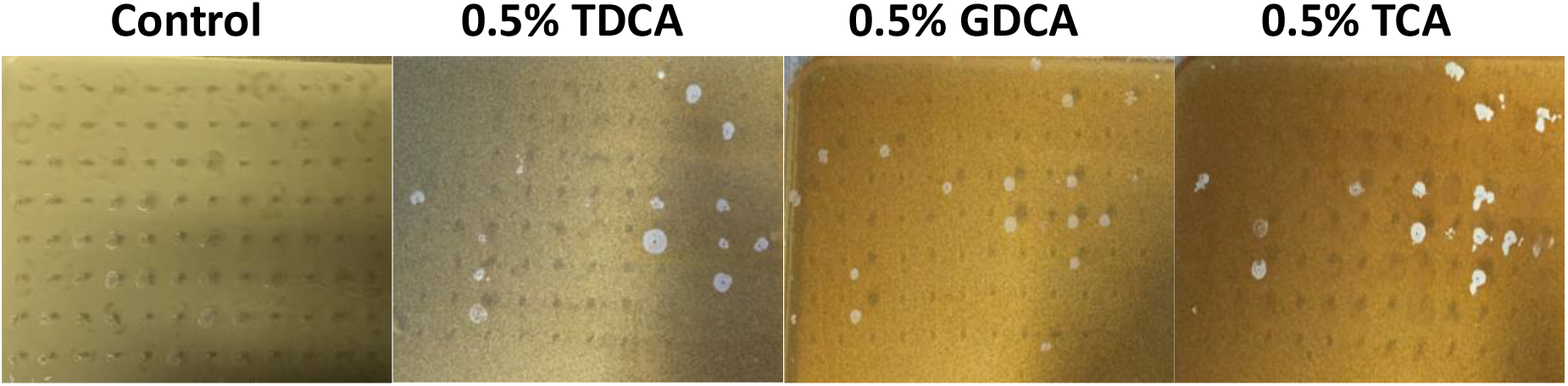
Detection of bile acid (BA) metabolising bacteria from formula fed infant faecal material. Assays were performed in the presence of conjugated BA’s deoxycholic acid (DCA) and cholic acid (CA). The presence of a white precipitate as seen above indicates that these representatives are potentially BSH active.

### Isolate distinction and species identification

RAPD analysis was performed as described by Ehrmann *et al*.,(Ehrmann *et al*., 2003) to the 60 BSH active isolates and benchmarked against a selection of *Lactobacillus* strains. A range of banding profiles were evident among the infant isolates which could be divided into four groups subsequently designated A-D as indicated in Figure 2. The majority of isolates formed two distinct groups (A and C) that were not similar, in profile, to any of the control strains. Group B showed a similar RAPD profile to *L. salivarius* UCC118 while group D had its own profile and was represented by one isolate. From RAPD analysis, the 60 BSH active infant isolates were represented as follows; Group A contained 38 (63%) representatives, there were 2 representatives (3%) for group B, 19 (32%) representatives for group C and 1 (1.5%) representative for group D.

**Figure 2:**
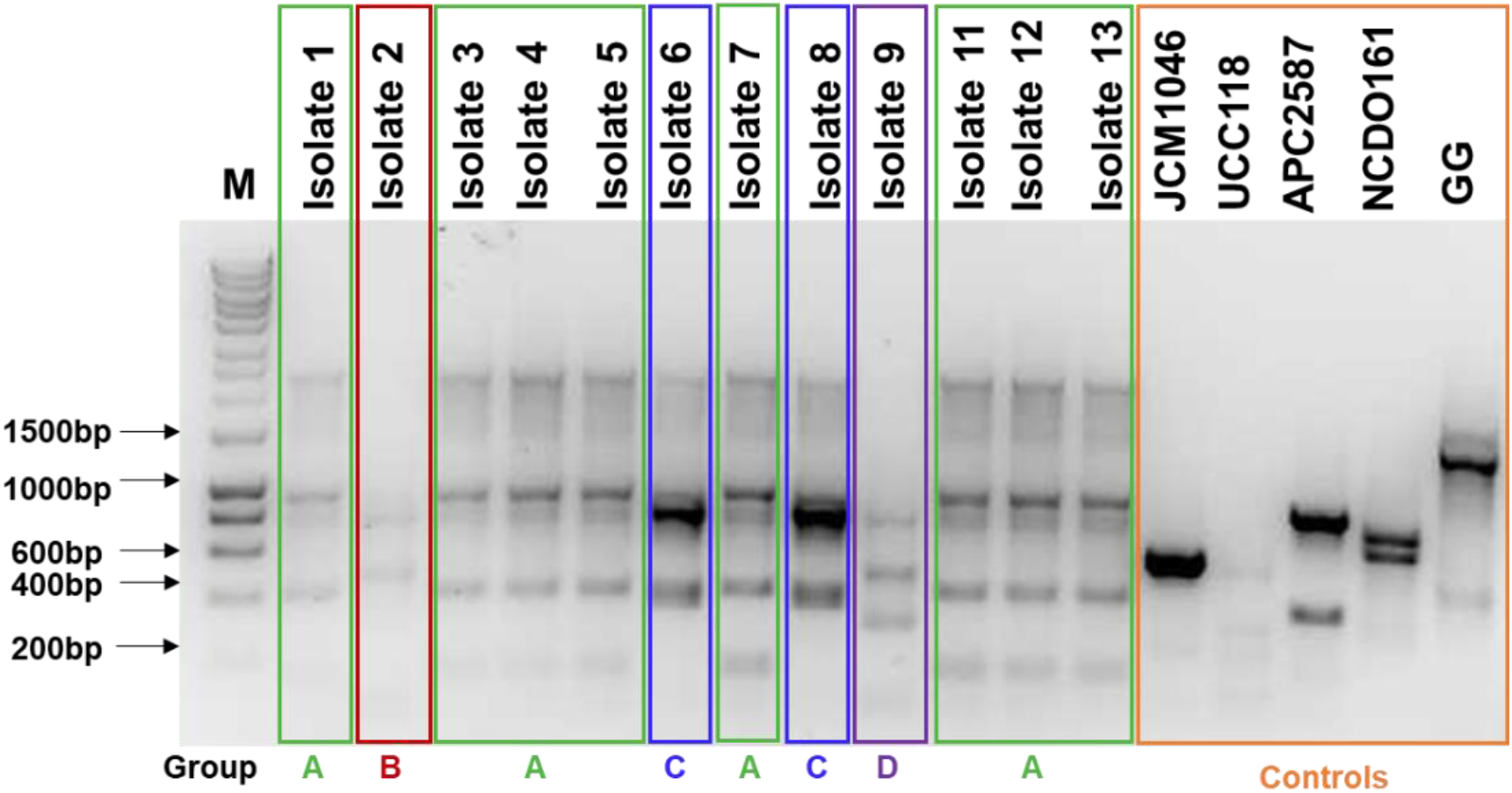
Representative Random Amplification of Polymorphic DNA (RAPD) analysis of BA metabolising infant isolates. Four main groups (A-D) were delimited as indicated. Control indicator strains are named as above: *L. salivarius* JCM1046, *L. salivarius* UCC118, *L. reuteri* APC2587, *L. casei* NCDO161 and *L. rhamnosus* GG.

16S rRNA genes from a number of representatives from each group of isolates were amplified using universal primers RW01 and DG74 (Greisen *et al*., 1994). Group A isolates shared 99% sequence identity to *Enterococcus durans*. One representative of this strain was selected for further study and designated APC2847. Variation was evident among group B isolates with one isolate displaying 99% sequence identity to *Enterococcus durans* while another isolate from the same group shared 99% sequence identity to *Enterococcus faecium*. Therefore, both of these isolates from Group B were selected for further study and they were designated APC2848 and APC2849 respectively. Group C isolates showed identical 16s rDNA sequence to each other and showed high levels of homology (99%) with *Lactobacillus acidophilus*. One of these strains, designated APC2845, was carried forward in this study. The single isolate in Group D, was 99% identical to *Streptococcus salivarius* and designated APC2846 but on further re-examination it was not BSH active. As a result, this strain was excluded from this study.

Multi locus sequence typing (MLST) analysis was performed for all isolates using a rage of different housekeeping genes. *Lactobacillus acidophilus* APC2845 species identification was confirmed following comparison with a number of characterised *L. acidophilus* strains (*L. acidophilus* NCFM, *L. acidophilus* ATCC314, *L. acidophilus* ATCC 4357 and *L. acidophilus* ATCC53544) based on three housekeeping genes: Translation elongation factor G (*fusA*), DNA gyrase subunit A (*gyrA*) and Recombinase A (*recA*)(Ramachandran *et al*., 2013)(Figure 3a). MLST analysis was also performed to delimit *Enterococcus* isolates APC2847, APC2848 and APC2849. Initial 16S rRNA sequencing identified APC2847 and APC2848 as *Enterococcus durans* and APC2849 as *Enterococcus faecium*. However, de novo genome sequencing pointed to all three of these isolates as potentially *Enterococcus casseliflavus* on the basis of complete genome sequence relatedness. Using the RAST annotated sequences of the housekeeping genes: ATP synthase subunit alpha (*atpA*), RNA polymerase subunit alpha (*rpoA*) and phenylalanine tRNA synthetase, alpha subunit (*pheS*), comparative genomics was performed against an array of different *Enterococcus* species which allowed clear differentiation and novel species detection(Naser *et al*., 2005a, Naser *et al*., 2005b). Phylogenetic trees generated using these sequences confirm, with a high degree of certainty, that all isolates cluster together with *Enterococcus avium* species contrary to original 16s rDNA sequencing (Figure 3b). *Enterococcus* sp. APC2847 clusters with *Enterococcus avium* LMG15118 while both *Enterococcus* sp. APC2848 and APC2849 clustered with *Enterococcus avium* LMG10744T. For further confirmation of this species assignment an *in vitro* assay was carried out to determine if these isolates could ferment the carbohydrate methyl-alpha-D-glucopyranoside (Devriese *et al*., 1996). All *Enterococcus* isolates were capable of producing acid from this carbohydrate which is a metabolic function frequently seen in *Enterococcus avium* species confirming the species identity of these isolates. For the remainder of this study these strains will be referred to as *Enterococcus avium* APC2847, *Enterococcus avium* APC2848 and *Enterococcus avium* APC2849.

**Figure 3:**
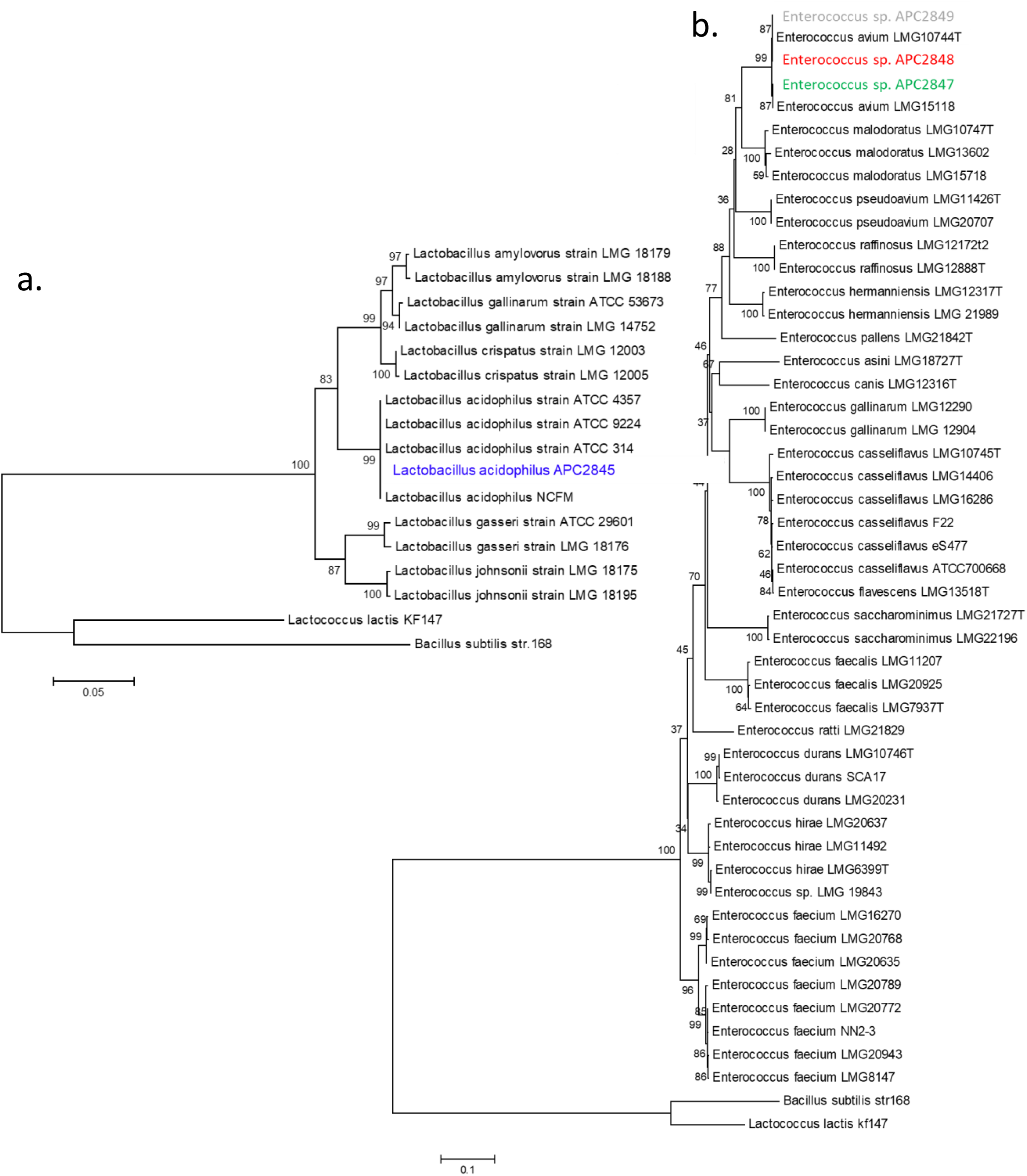
Neighbor-joining tree for species identification of isolates based on a) fusA gene sequences. *Lactobacillus acidophilus* APC2845 was compared to 14 *Lactobacillus* strains belonging to the *L. acidophilus* complex. *L. acidophilus* APC2845 clustered together with other *L. acidophilus* strains and b) atpA gene sequences. *Enterococcus* isolates APC2847, APC2848 and APC2849 were compared to 44 Enterococcal strains of various species where it was evident that these isolates cluster together with the species *E. avium*. Bootstrap values after 1,000 repetitions are indicated with *Lactococcus lactis* and *Bacillus subtilis* included as outgroups.

### Evaluation of the presence of bile salt hydrolase genes in infant isolates *L. acidophilus* APC2845, *E. avium* APC2847, *E. avium* APC2848 and *E. avium* APC2849

*L. acidophilus* APC2845, representing 32% (19/60) of infant bile deconjugating isolates, contained two putative *bsh* genes designated *bsh1* and *bsh2*. Both genes are 978bp in length encoding two distinct 325 amino acid proteins which are 58% identical to each other on the basis of amino acid comparison. Relative to other *L. acidophilus* BSH proteins, BSH1 is highly similar to that reported for *L. acidophilus* NCFM BSHA (100%), *L. acidophilus* LA4 BSHA (99%) and *L. acidophilus* LA11 BSHA (100%)(accession numbers: CP000033, ACL98173.1, ACL98175.1 respectively), with BSH2 showing high degrees of similarity to *L. acidophilus* NCFM BSHB (100%), *L. acidophilus* LA4 BSHB (100%) and *L. acidophilus* LA11 BSHB (99%) (Accession numbers: CP000033, ACL98174.1, ACL98176.1 respectively)(Figure 4a). Comparative analysis confirms that both BSHs carried by *L. acidophilus* APC2845 are 100% identical to *L. acidophilus* NCFM on the basis of amino acid sequences. Examination of the proposed conserved amino acid residues required for catalytic function of BSH enzymes were examined. These residues include: Cys2, Arg16, Asp19, Asn79, Asn171 and Arg224 (Fang *et al*., 2009a) (Fang *et al*., 2009b) and are present in both BSH1 and BSH2 proteins in isolate *L. acidophilus* APC2845 indicating that both of these enzymes may contain a full, intact, active site and therefore demonstrate BSH activity.

**Figure 4:**
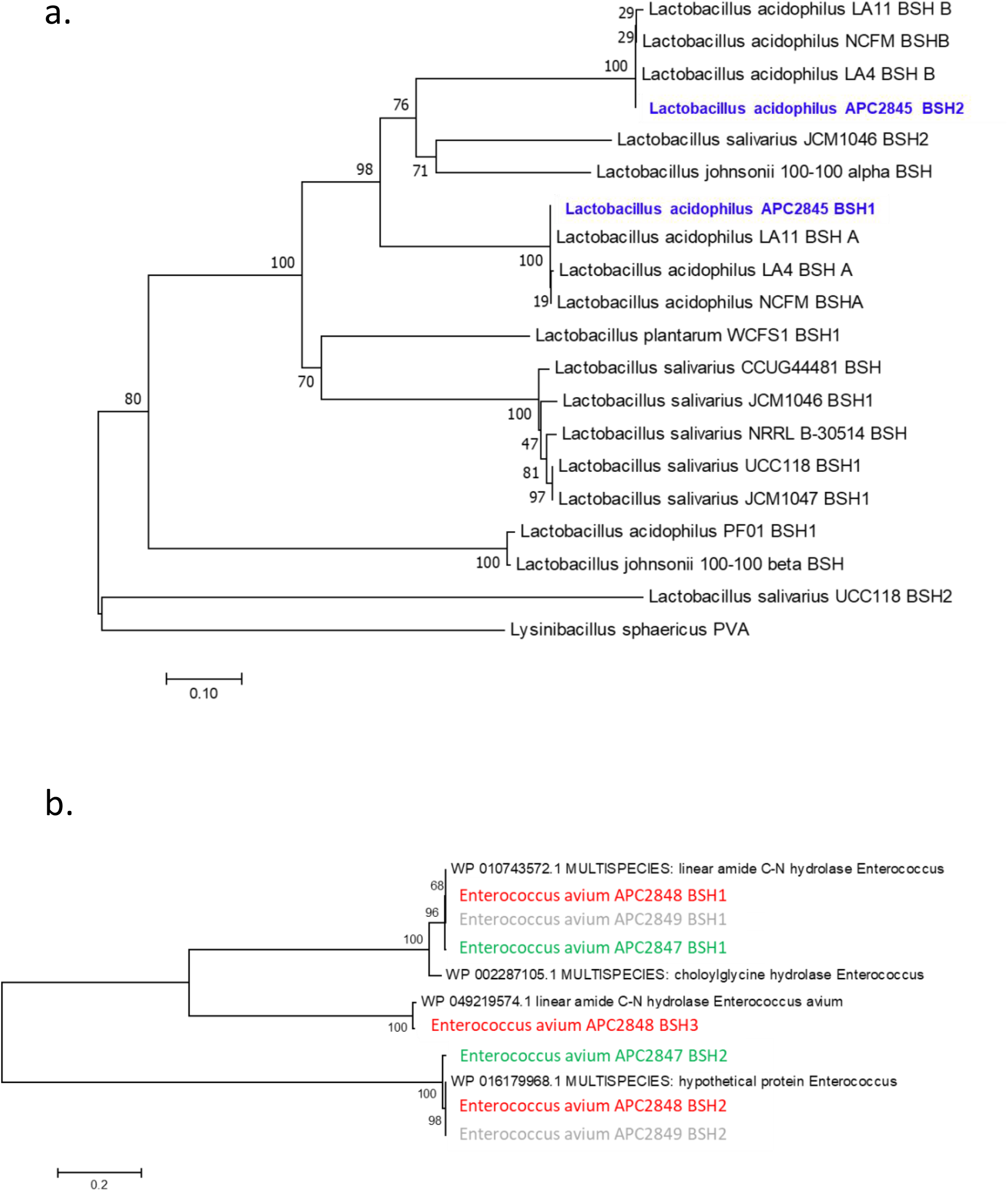
Phylogenetic analysis of BSH proteins. a) *L. acidophilus* APC2845 BSH1 and BSH2 amino acid sequences. Comparisons are made against BSH from different *Lactobacillus* species, and b) *E. avium* APC2847, APC2848, APC2849 BSH amino acid sequences. The BSH sequences for each of these isolates were compared to annotated BSHs from deposited *Enterococcus* species and *Enterococcus avium* ATCC14025. Multiple sequence alignment and phylogenetic tree construction was performed by MEGA6 software(Tamura *et al*., 2013). *Lysinibacillus sphaericus (L. sphaericus* LMG22257 PVA) was included as outgroup.

*Enterococcus avium* APC2847 (group A, representing 64% of bile acid deconjugators) revealed two possible *bsh* genes, designated *bsh1* and *bsh2*, which could be annotated as choloylglycine hydrolase (EC 3.5.1.24). *bsh1* is 975bp in length and encodes a 324 amino acid protein whereas *bsh2* is 1008bp encoding a 335 amino acid protein. These sequences were collectively analysed further with *Enterococcus avium* APC2848 and 2849 sequences below for identity and conservation of residues as well as potential functionality. *Enterococcus avium* APC2848 (group B-collectively with APC2849 representing 3.5% of infant bile acid deconjugators) contained three possible *bsh alleles*, that were subsequently designated *bsh1, bsh2* and *bsh3. bsh1* was 975bp in length encoding a 324 amino acid protein with *bsh2* composed of 1008bp encoding a 335 amino acid protein. Finally, a third *bsh (bsh3)* of 891bp encoding a 296 amino acid protein was identified within this isolate. *Enterococcus avium* APC2849 also contained two possible *bsh* genes which were 100% identical to their respective proteins identified in group B isolate *Enterococcus avium* APC2848.

Deposited BSH sequences were sought in order to analyse the infant *Enterococcus avium* isolates. Only six *Enterococcus* BSH amino acid sequences were returned, genbank accession numbers: WP010743572.1, WP016179968.1, WP002287105.1 and one *Enterococcus avium* ATCC14025 BSH amino acid sequence genbank accession number: WP049219574.1. These data were subsequently applied for analysis and comparison with the infant isolated *Enterococcus avium* APC2847, APC2848 and APC2849 BSH amino acid sequences. Although numerous BSH protein sequences from different species of *Enterococcus* such as *E. faecalis* have been deposited to Genbank, only sequences (accession numbers mentioned above) which show high % homology to *E. avium* BSH were selected for comparative analysis. Amino acid sequence alignment and phylogenetic trees confirm that BSH1 from *E. avium* APC2847, APC2848 and APC2849 cluster together with *Enterococcus* multispecies linear amide C-N hydrolase (WP010743572.1) and the *Enterococcus* multispecies choloylglycine hydrolase (WP002287105.1). *E. avium* APC2847 showed 99% homology to WP010743572.1, and 93% sequence identity to WP002287105.1. *E. avium* APC2848 and APC2849 shared 100 % homology to WP010743572.1 closely followed by 93% sequence identity to WP002287105.1 (Figure 4b). BSH2 sequences also clustered together in a distinct group with *E. avium* APC2847 and both *E. avium* APC2848 and APC2849 sharing 99% and 100% identity, respectively to an *Enterococcus* multispecies hypothetical protein (WP016179968.1). *E. avium* APC2848 BSH3 was the most divergent BSH sequence of the newly identified isolates and showed the highest homology (99%) with the only other *E. avium* BSH sequence included in this analysis (*Enterococcus avium* linear amide C-N hydrolase WP049219574.1). BSH1 proteins identified in *E. avium* APC2847, AP2848 and APC2849 contain all of the critical residues (Cys2, Arg16, Asp19, Asn79, Asn170 and Arg223) at the proposed locations and distances indicating that these enzymes should contain full intact active sites. In contrast, at these critical active site positions, severe alterations Tyr2, Lys16, Met19, Val79, Val170 and Val223 for BSH2 and Ile2, Gln16, Lys19, Glu79, Asn170 and Phe223 for BSH3 were evident and indicate that they may not function in bile acid metabolism.

### All infant isolates, *L. acidophilus* APC2845, *E. avium* APC2847, *E. avium* APC2848 and *E. avium* APC2849 collectively display *in vitro* bile altering capabilities which can be attributed to one or more individual BSH proteins contained with each isolate

In order to determine the specific BA metabolism ability of isolates *L. acidophilus* APC2845, *E. avium* APC2847, *E. avium* APC2848 and *E. avium* APC2849, co-incubation assays (n=3) with these isolates were performed in 0.5% porcine (see materials and methods) against a selection of known BSH active strains (*L. salivarius* JCM1046 and *L. reuteri* APC2587) and BSH inactive strains (*L. salivarius* UCC118, *L. casei* NCDO161 and *L. rhamnosus* GG). It is evident that all infant isolates and benchmark strains, that carry BSHs, are metabolising bile acid when compared to non BSH carrying strains and to media containing bile acid alone. Here, lower levels of conjugated bile acids are evident with higher levels of primary and secondary bile acid classes (Figure 5a-c). *L. acidophilus* APC2845 and *E. avium* APC2848, consistently show relatively weak alterations.

**Figure 5:**
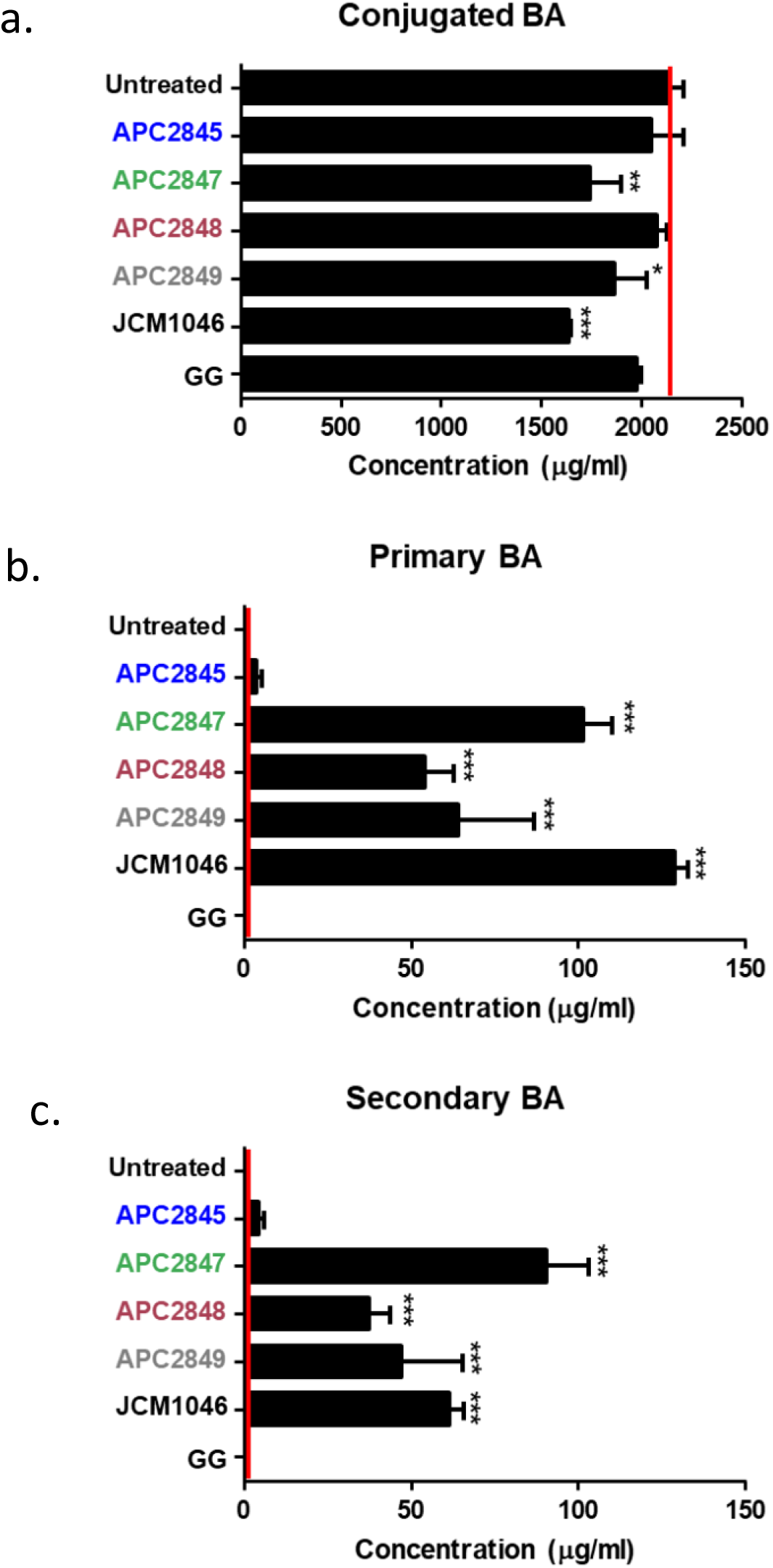
Assessment of deconjugating ability of infant isolates. A reduction in a) conjugated BAs with a corresponding accumulation of b) primary and c) secondary BAs can be seen in all BSH containing isolates. Bile acid concentrations were quantified against standard curves for each analyte and normalized according to internal standards using WatersR Targetlynx software. Data was plotted using GraphPad Prism v5.0 and is represented as mean ± standard deviation (SD). Statistical significance is indicated and calculated using one-way ANOVA followed by Dunnett’s post-hoc test: *\*= p<0.05; **=p<0.01; ***=p<0.001*.

In order to determine what individual enzymes are contributing the BSH activity displayed, each *bsh* gene present from each isolate were cloned into the pET-21b^+^ vector and expressed in *E. coli* BL21 (DE3) cells. The co-incubation assay was repeated as before using the pET-21b^+^ empty vector as a negative control. Both BSH1 and BSH2 proteins from *L. acidophilus* APC2845 appear to be active through reducing concentrations of conjugated bile acids (Figure 6a) which lead to an accumulation of freed primary and secondary BAs (Figure 6b and 6c respectively). Both proteins result in the liberation of free CA, CDCA, DCA, LCA, UDCA, HCA, HDCA, α/ω-MCA, β-MCA, however, heatmap analysis revealed the extent of the substrate specificity for each BSH protein with BSH1 showing preference for glyco conjugated BAs while BSH2 is capable of metabolising a range of both glyco and tauro conjugated moieties (Table 1). On the other hand, BSH activity displayed by *E. avium* APC2847, *E. avium* APC2848 and *E. avium* APC2849 appear to be solely demonstrated by BSH1 proteins. These BSH1 proteins also lead to the accumulation of CA, CDCA, DCA, LCA, UDCA, HCA, HDCA, α/ω-MCA, β-MCA through the deconjugation of a wide range of glyco and tauro conjugated BAs. No activity was seen for APC2847-BSH2, APC2848-BSH2 and APC2848-BSH3. Overall these results appear to confirm the *in silico* analysis carried out one each individual BSH protein.

**Figure 6:**
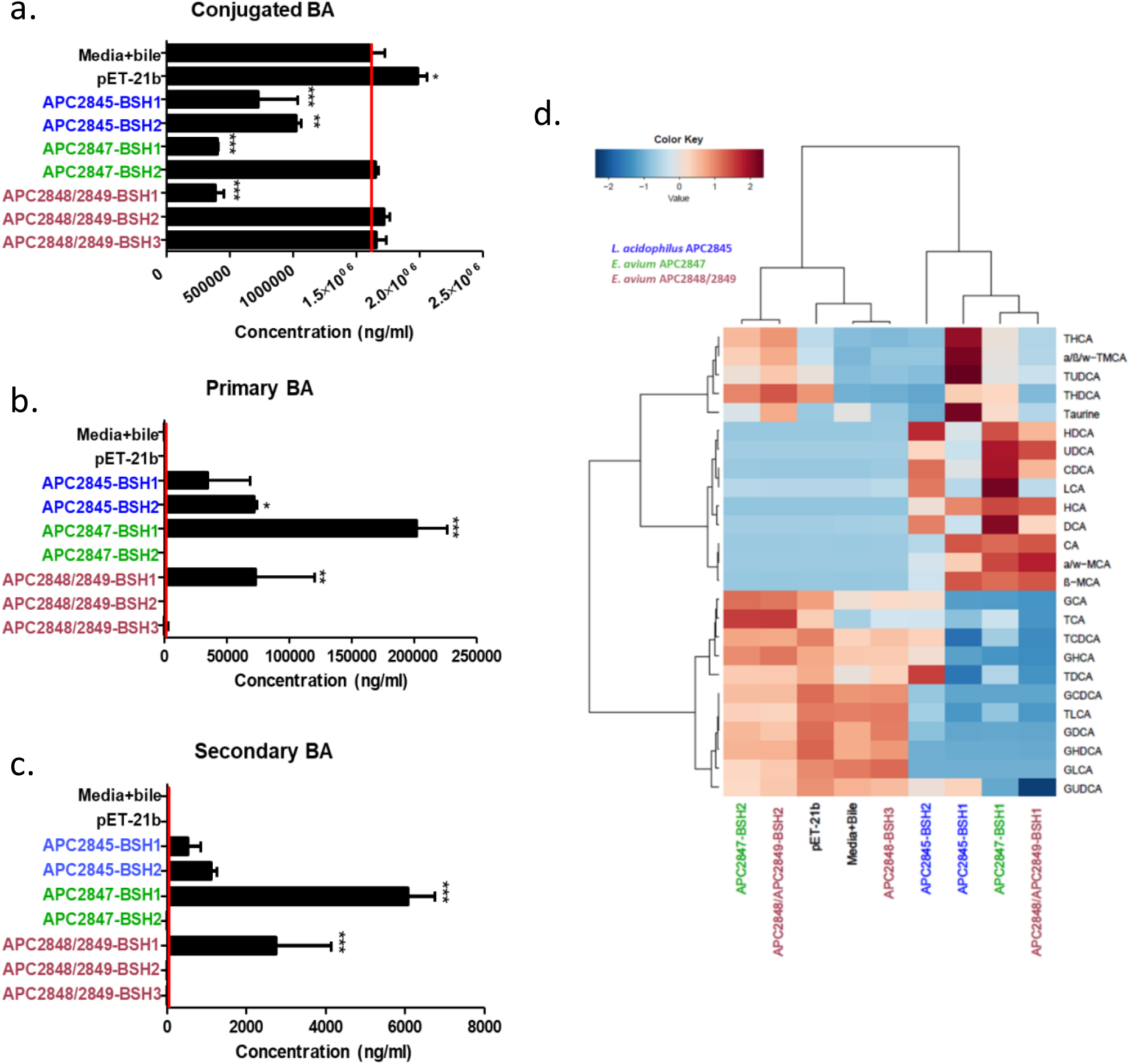
*L. acidophilus* APC2845, *E. avium* APC2847, APC2848 and APC2849 individual BSH protein activity. a) Lower conjugated BAs with a corresponding accumulation of b) primary and c) secondary BAs indicated which BSH proteins are active. d) Heatmap demonstrate the range of bile acid moieties metabolised by both BSH1 and BSH2 proteins. Bile acid concentrations were quantified against standard curves for each analyte and normalized according to internal standards using WatersR Targetlynx software. Data was plotted using GraphPad Prism v5.0 and is represented as mean ± standard deviation (SD). Statistical significance is indicated and calculated using one-way ANOVA followed by Dunnett’s post-hoc test: *\*= p<0.05; **=p<0.01; ***=p<0.001*. Heatmap was generated using RStudio software version 1.1.453.

**Table 1:** Summary of substrate specificity for each individual BSH protein isolated from strains *L. acidophilus* APC2845, *E. avium* APC2847, APC2848 and APC2849.

| Strain | Glyco-conjugates | Tauro-conjugates | Accumulated Bile Acids |
| --- | --- | --- | --- |
| *L. acidophilus* APC2845 BSH1 | GCA, GCDCA, GDCA, GLCA, GUDCA, GHCA, GHDCA | TCDCA, TDCA, TLCA | CA, CDCA, DCA, LCA, UDCA, HCA, HDCA, $\alpha/\omega$-MCA, β-MCA |
| *L. acidophilus* APC2845 BSH2 | GCDCA, GDCA, GLCA, GUDCA, GHCA, GHDCA | TLCA | CA, CDCA, DCA, LCA, UDCA, HCA, HDCA, $\alpha/\omega$-MCA, β-MCA |
| *E. avium* APC2847 BSH1 | GCA, GCDCA, GDCA, GLCA, GUDCA, GHCA, GHDCA | TCDCA, TDCA, TLCA | CA, CDCA, DCA, LCA, UDCA, HCA, HDCA, $\alpha/\omega$-MCA, β-MCA |
| *E. avium* APC2847 BSH2 | No activity | No activity | No activity |
| *E. avium* APC2848/APC2849 BSH1 | GCA, GCDCA, GDCA, GLCA, GUDCA, GHCA, GHDCA | TCA, TCDCA, TDCA, TLCA | CA, CDCA, DCA, LCA, UDCA, HCA, HDCA, $\alpha/\omega$-MCA, β-MCA |
| *E. avium* APC2848/APC2849 BSH2 | No activity | No activity | No activity |
| *E. avium* APC2848/APC2849 BSH3 | No activity | No activity | No activity |

### Assessment of *L. acidophilus* APC2845, *E. avium* APC2847, *E. avium* APC2848 and *E. avium* APC2849 tolerance to porcine bile

A prerequisite for potential probiotic strains is their ability to resist the toxic effects of bile (FAO/WHO 2002, IPA 2017) ENREF 8. In order to determine if the infant isolated *L. acidophilus* APC2845, *E. avium* APC2847, *E. avium* APC2848 and *E. avium* APC2849 could tolerate the effects of bile, these isolates were exposed to 0.5% porcine bile and growth was monitored over a 24-hour period (Figure 7). All isolates examined were capable of growth in the MRS media with and without porcine bile. *L. acidophilus* APC2845 grew optimally in 0.5% porcine bile (Figure 7d). Similar to that seen with *L. acidophilus* APC2845, isolates *E. avium* APC2847, *E. avium* APC2848 and *E. avium* APC2849 were delayed for growth in MRS alone compared to bile supplemented media. In summary, all strains display high levels of bile tolerance when exposed to glyco dominant porcine bile.

**Figure 7:**
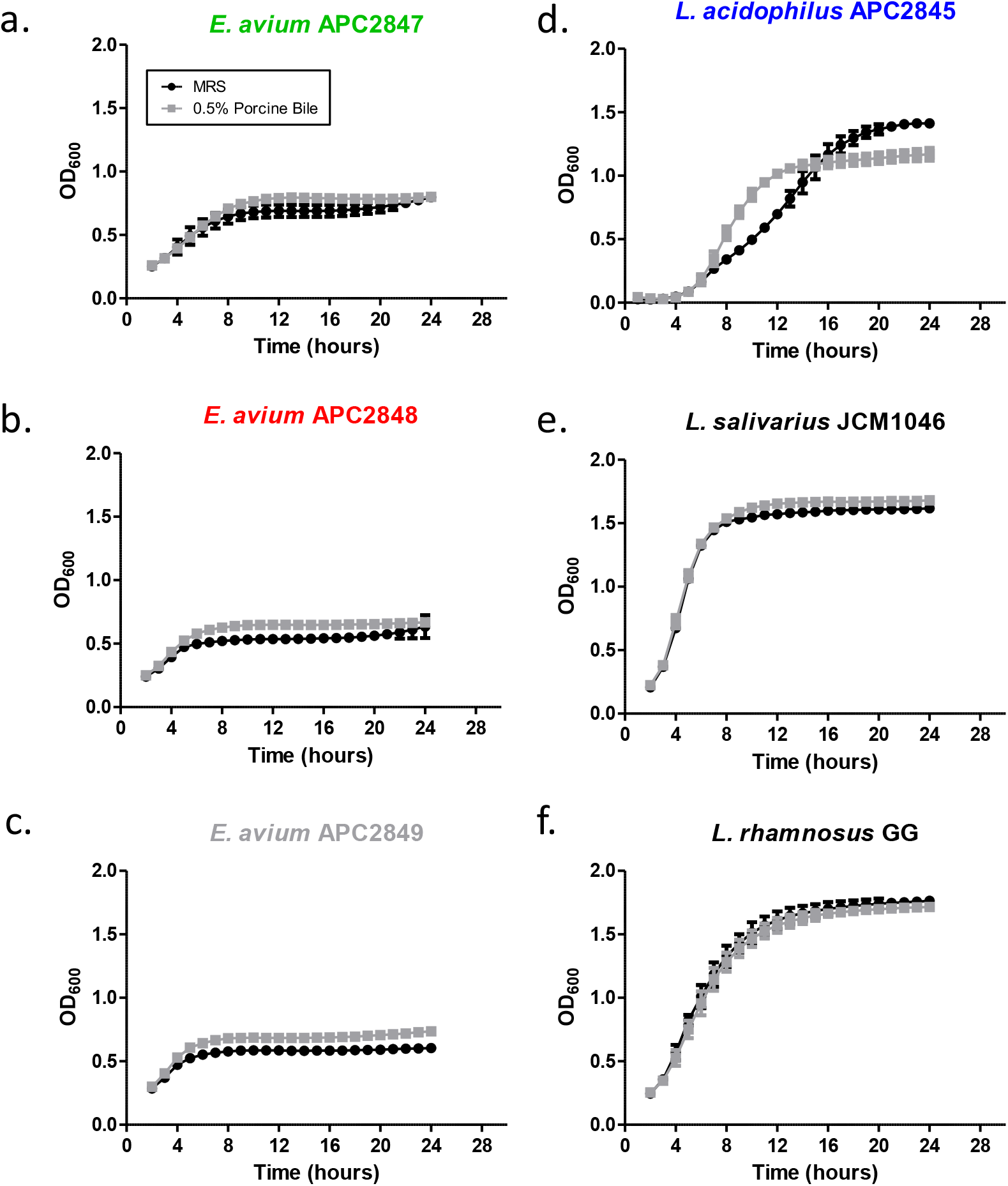
Bile tolerance of infant isolates and benchmark strains in the presence of 0.5% porcine bile. Growth assessment of (a) *E. avium* APC2847, (b) *E. avium* APC2848, (c) *E. avium* APC2849 (d) *L. acidophilus* APC2845, (e) *L. salivarius* JCM1046 and (f) *L. rhamnosus* GG. Error bars represent the mean ± standard deviation from the mean (SD) in triplicate experiments (n = 3).

### Survival and benchmarking of *L. acidophilus* APC2845, *E. avium* APC2847, *E. avium* APC2848 and *E. avium* APC2849 on exposure to simulated gastrointestinal conditions

All of the isolates tested demonstrated the ability to survive passage through the GIT model (Figure 8). Full transit though the simulated GIT resulted in a net loss of cell viability of approximately 1.23 log_10_ fold for *L. acidophilus* APC2845, 0.89 log_10_ fold for *E. avium* APC2847, 2.27 log_10_ fold for *E. avium* APC2848 and 0.78 log_10_ fold for *E. avium* APC2849. Comparing these to the benchmark strains: *L. salivarius* JCM1046 displayed a 1.93 log_10_ fold decrease in cell viability with *L. rhamnosus* GG showing a 0.18 log_10_ fold decrease. Overall each strain varied in their tolerance and viability at the different points of passage, pH and exposure to digestive enzymes through the simulated GIT. Overall *E. avium* APC2849 and *E. avium* APC2847 showed the highest tolerance and survival throughout the GIT. This was followed by *L. acidophilus* APC2845 and *E. avium* APC2848. These results are particularly interesting with respects to the three *E. avium* isolates which contain different numbers of BSH genes and varying levels of BSH activity, however it is important to note that these effects may be independent of BSH activity and may be a feature of other encoded properties of these new isolates.

**Figure 8:**
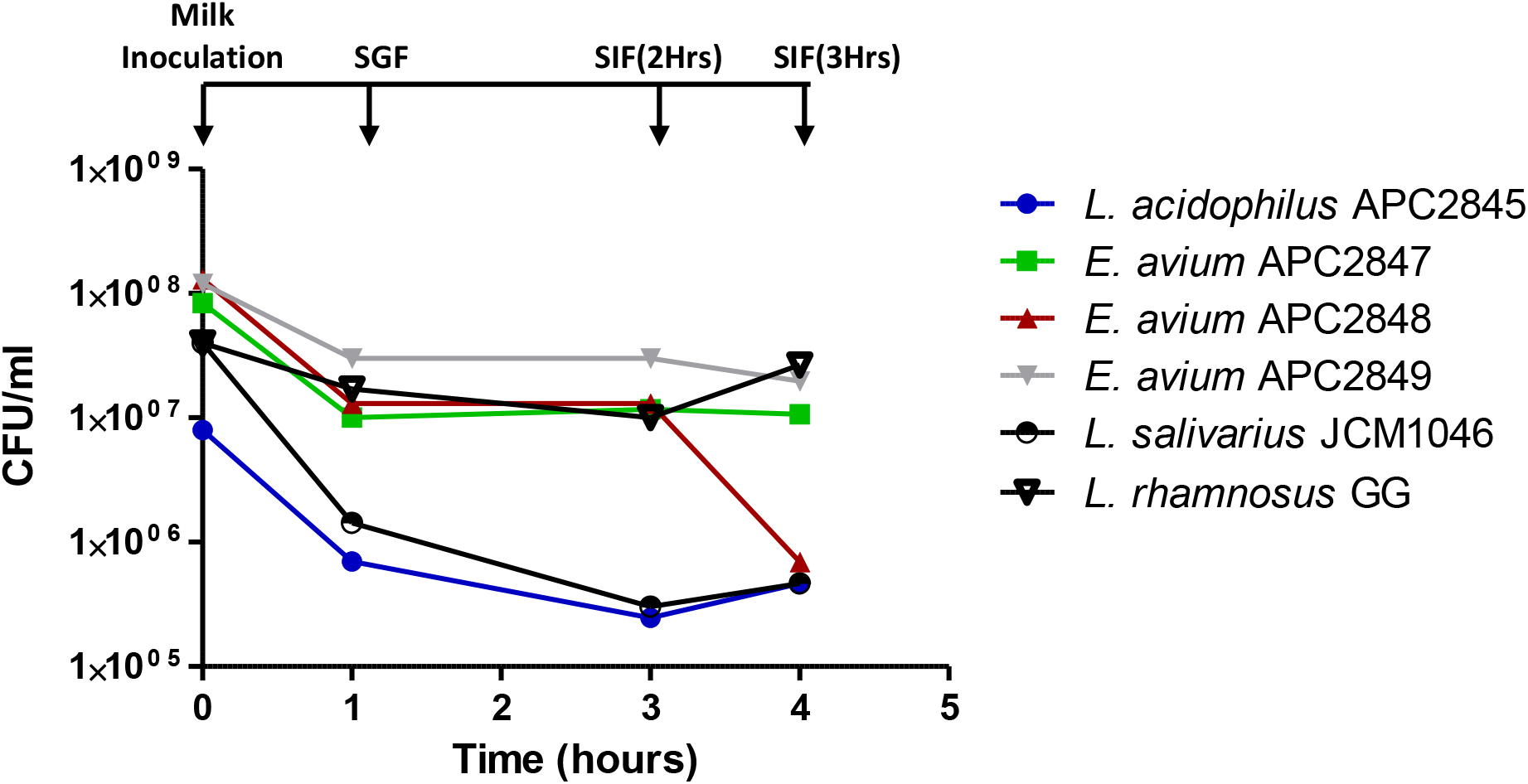
Survival of infant isolates and benchmark strains through the simulated gastrointestinal tract (GIT). Graph details the effects on viability of *L. acidophilus* APC2845 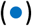, *E. avium* APC2847 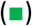, *E. avium* APC2848 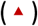, *E. avium* APC2849 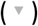, *L. salivarius* JCM1046 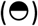 and *L. rhamnosus* GG (∇). Arrows indicate addition of simulated gastric juice (SGF) (1-hour) and simulated intestinal fluid (SIF) (after 2 and 3 hours). Bacterial colony forming units (CFUs) were calculated post assessment on solid media. Error bars represent the mean ± standard deviation from the mean (SD) of 3 biological replicates (n = 3).

## Discussion

Studies based on functionality of the infant gut are lacking and this study aimed to examine a specific important microbial gut function, bile acid metabolism. An infant faecal sample was chosen for this study, since in this 0-2 years period, rapid growth and metabolism is critical and bile acids are recognised as signalling hormones, they have a central role in metabolism, particularly lipid metabolism, and weight gain, they are also good indicators of microbial change. Bile acids can also influence human health through their intrinsic properties (Enright *et al*., 2017) and also by their interactions with different receptors both locally and systemically (Long *et al*., 2017b).

This study isolated and characterised BSH representative bacterial isolates of infant origin. From a bile agar assay screen of 792 isolates, less than 10% of detected microbes, 60 bacterial isolates, proved capable of deconjugating key receptor activating bile acids (BAs) GDCA, TDCA and TCA. Altered BA metabolism may be an adaptation of the infant gut to sustain micellar formation and nutrient absorption, particularly lipid and its fat-soluble vitamins D, E, K, A, for further rapid metabolism.

Of these BA metabolising isolates RAPD analysis identified four distinct groups (named A, B, C, D). Group C or 32% of BA metabolising isolates was represented by *Lactobacillus acidophilus* species and groups (A, B) or 65% of BA metabolising isolates were identified as *Enterococcal* species *avium* through a combination of 16S analysis, newly developed MLST systems, genomic comparisons and by phenotypic acidification of methyl-a-D-glucopyranoside.

*Lactobacillus acidophilus* APC2845 was identified through a combination of 16S rDNA (99% identity) and MLST. Genetic diversity within this species is quite low (Bull *et al*., 2014) but as expected, APC2845 clustered together with all *L. acidophilus* strains compared in this study. *L. acidophilus* APC2845 contained two distinct *bsh* coding genes designated *bsh*1 and *bsh*2, which showed 100% identity to those designated *bshA* and *bshB* in *L. acidophilus* NCFM (McAuliffe *et al*., 2005). *L. acidophilus* APC2845 activity was analysed by UPLC TMS and exact quantity modifications were assessed in *in vitro* bacteria and bile co-incubation assays. *L. acidophilus* APC2845 showed a restricted range of conjugated substrate preference for both glyco and tauro-conjugated bile salts when compared to other benchmark *Lactobacillus* species where both BSH proteins it contained contributing to the effects seen. For other *L. acidophilus* strains; *L. acidophilus* NCFM (2 BSH enzymes), both is active against some glyco and tauro BAs and, its BSH2 lacks activity against tauro-conjugated moieties (McAuliffe *et al*., 2005). *L. acidophilus* LA11 is active towards TCA, TDCA, TCDCA, GCA, GDCA with *L. acidophilus* LA4 showing decreased activity towards GCDCA, TCA, TDCA and TCDCA when examined using a modified ninhydrin assay protocol (Jiang *et al*., 2010). *L acidophilus* APC2845 could survive oral, gastric and intestinal phase transit with the total net loss of only 1.5 log fold, performing well against *Lactobacillus* benchmark strains. When exposure, growth and tolerance of porcine bile acid were conducted independently this strain also displayed the ability to grow well in the presence of bile. Given the low genetic diversity among the *L. acidophilus* species there is a debate as to whether it is possible to assume and assign the same functionality to all strains in this monophyletic species (Bull *et al*., 2014). The results described herein, the clonality of *L. acidophilus* strains, their long history of dairy application and their <u>Q</u>ualified <u>P</u>resumption of <u>S</u>afety (QPS) status (EFSA 2007, Rychen *et al*., 2017) would further solidify the application of *L. acidophilus* APC2845 as a probiotic for human consumption.

The three *Enterococcus avium* isolates identified in the infant faecal sample are also classified as Lactic acid bacteria (LAB). Enterococci are commensal strain but certain members of the species can also be pathogenic leading to questions of safety when applied as probiotics and because of this *Enterococcus* have not been awarded QPS status requiring in depth characterisation of any new isolates (Gueimonde *et al*., 2004, Huys *et al*., 2013, Ricci *et al*., 2017). *Enterococcus* spp. can frequently be found in foods such as cheese, sausage and olives (Foulquié Moreno *et al*., 2006) with *E. faecium* and *E. faecalis* the only *Enterococcus* species applied as probiotics (Franz *et al*., 2011, Celiberto *et al*., 2017). Many studies report the potential beneficial effects of other *Enterococcus* species such as *E. durans* M4-5 which demonstrates an anti-inflammatory effect through the production of the SCFA butyrate (Avram-Hananel *et al*., 2010) and strains of *E. hirae* which are capable of modulating and facilitating the effects of anti-tumorigenic drugs (Arokiyaraj *et al*., 2014, Gupta and Tiwari 2015, Daillère *et al*., 2016). Confirmation of the identity of these three distinct *Enterococcus avium* isolates APC2847, APC2848 and APC2849 was identified through a combination of genetic (16S rDNA and MLST analysis) and phenotypic assays (acidification of methyl-a-D-glucopyranoside). Interestingly, *Enterococcal* species are often pathogenic in nature, and are often associated with bloodstream and urinary tract infections as well as intra-abdominal infections, meningitis and endocarditis(Na *et al*., 2012b). However, infections can be fatal in over 30% of cases where an underlying condition already exist (Patterson *et al*., 1995). In a study of 2,672 Enterococcus-positive bacteraemia cases *E. avium* infection represented just 2.3% of cases (Na *et al*., 2012a). These authors showed that bacteraemia was due to polyclonal infections where *E. avium* was associated with the presence of species in particular *E. coli*. All patients presented with sepsis and mortality was 25%. Interestingly, the underlying disease for 58% of these patients was biliary disease.

*Enterococcus avium* APC2847, APC2848 and APC2849 were further characterised according to their BSH activity. This is the first report on bile salt metabolism ability by these species. BSH activity has been reported in a number of *Enterococcus spp*. including *E. faecalis* (Chand *et al*., 2016, Chand *et al*., 2018), *E. faecium, E. durans, E. gallinarum* and *E. casseliflavus* (Franz *et al*., 2001), however BSH activity for the species *E. avium* has not been reported in the literature. *E. avium* APC2847 contained two distinct *bsh* coding genes designated *bsh1* and *bsh2. E. avium* APC2848 shares high homology with *E. avium* APC2849 with both strains containing identical *bsh* coding genes also designated *bsh*1 and *bsh*2 with *E. avium* APC2848 differing from APC2849 by the presence of a third *bsh* gene (*bsh*3). Blast analysis revealed that all BSH1 protein had high sequence identity to linear C-N amide hydrolase found within multi-species *Enterococcus*. In contrast, BSH2 and BSH3 proteins are unlikely to be active given that the former showed high homology to a hypothetical protein and the later showed high similarity to linear C-N amide (Chand *et al*., 2016, Chand *et al*., 2018). In support of this analysis of active site residues for these BSHs were considerably altered, also these proteins lacked the autocatalytic Cys2 residue which is crucial for activity (Brannigan *et al*., 1995, Xu *et al*., 2016, Chand *et al*., 2018).

The bile acid metabolizing ability of *Enterococcus avium* APC2847, APC2848 and APC2849 was also collectively and individually examined using UPLC-MS. Analysis confirmed that all of these strains have the capacity to deconjugate BAs and that BSH1 proteins only are responsible for the activity displayed by these isolates. Distinct differences in the levels of activity were also evident across the strains with *E. avium* APC2847-BSH1 displayed the highest activity followed by *E. avium* APC2848/2849-BSH1. All three *E. avium* strains demonstrated the ability to survive exposure to 0.5% porcine bile and survive transit through the simulated GIT, however, due to the rare incidence of *E. avium* infection and bacteraemia (Patel *et al*., 1993, Na *et al*., 2012a), it is unlikely that any of these three *E. avium* infant isolates would be recommended for use as probiotics (Montealegre *et al*., 2016, Bonacina *et al*., 2017, Jung *et al*., 2017). However, this warrants further investigation into the role of these species in infant gut development and maturation given the fact that these strains are a feature of formula fed infants

## Methods

### Preparation of infant bank of bacteria isolates

Ethics were granted under InfantMet project ref ECM 3(w) 07/02/12. One individual, fresh faecal sample was obtained from a 2 year old female infant donor with no previous exposure to antibiotics for at least 6 months prior to giving sample. Sufficient sterile PBS was added to a concentration of 250 mg/ml. The mixture was homogenized, then filtered through a 70μm cell strainer (BD) before serial dilution in sterile phosphate buffered saline (PBS, Sigma) then spread onto MRS agar (Oxoid). These agar plates were incubated under anaerobic conditions at 37°C for 48 hours to facilitate growth. Single colonies were selected for anaerobic growth in MRS broth (Oxoid) in deep 96-well plates. In total 792 individual isolates were stocked and stored at −80°C in 15% glycerol for further analysis

### Bacterial Strains, Media and culture conditions

For the most part, bacteria were grown in the relevant liquid and agar media-Man Rogosa Sharpe (MRS) for *Lactobacillus* species and *Enterococcal* species at 37°C unless otherwise stated. Strains were grown either anaerobically (placed in an anaerobic chamber) or aerobically (by agitation) over 24-48 hours. Isolates were maintained as a frozen stock at −80°C in 15 % (v/v) glycerol. A selection of bile salt hydrolase genes were cloned independently into the vector pET-21b^+^ under the expression of the T7 promoter. These constructs were synthesised by Genscript (Netherlands) and expressed in *Escherichia coli* BL21 (DE3) cells (Invitrogen). *E. coli* strains were grown in Luria–Bertani (LB) broth at 37°^C^ with ampicillin (50μg/ml).

### Functional selection of bile metabolising infant isolates using bile agar assay

MRS agar (Oxoid) was prepared containing individual bile salts (0.5% w/v; GDCA, TDCA, TCA, Sigma), and CaCl2 (0.37 5 g/l, Sigma). Each bile salt was tested independently and MRS agar with no bile salts present was used as a negative control for comparison. Fresh overnight cultures of bacterial isolates, prepared as described above, were inoculated into punctures in MRS agar. These agar plates, containing the different isolates, were incubated at 37°C for 72 hours under anaerobic conditions. Any isolate positive for BSH activity either presented as a colony morphology change and /or the presence of a visible halo or precipitate of deconjugated and therefore hydrophobic bile salt form.

### Isolate distinction, species identification and genome analysis

Genomic DNA was purified using GenElute Bacterial Genomic DNA Kit (Sigma) according to the manufacturer’s guidelines. To distinguish differences in isolates Random Amplification of Polymorphic DNA (RAPD) PCR was carried out as described by Ehrmann *et al*., (2003)(Ehrmann *et al*., 2003). 16s rRNA sequencing was carried out to that described by Greisen *et al*., 1994 (*Greisen *et al*., 1994*) and *de novo* sequencing was carried out by MicrobesNG (https://microbesng.uk/). Predicted protein-coding genes within the graft genome sequencing reads were annotated by the RAST Annotation Server (Aziz *et al*., 2008). MEGA software version 6.0(Tamura *et al*., 2013) (http://www.megasoftware.net/) was used to perform multiple sequence alignments and phylogenetic trees. Nucleotide and amino acid sequences were analysed using the BLAST2 program (NCBI) and the CLUSTALW software package (Aiyar 2000). *Lactobacillus acidophilus* MLST was performed according to Ramachandran *et al*., (2013)(Ramachandran *et al*., 2013) while MLST for *Enterococcus* identification was carried out as described by Naser *et al*.,(2005)(Naser *et al*., 2005a, Naser *et al*., 2005b). Further phenotypic confirmation required for *Enterococcus* isolates was performed using a method developed by Devrise *et al*.,(1996)(Devriese *et al*., 1996) through the acidification of Methyl-α-D-Glucopyranoside.

### Assessment of bile acid tolerance and survival through a simulated model of the gastrointestinal tract of isolates: *L. acidophilus* APC2845, *E. avium* APC2847, *E. avium* APC2848 and *E. avium* APC2849

Growth assessment was performed in the presence and absence of bile isolated from porcine gall bladders. Strains were normalised 0.1 OD_600nm_ and were grown at 37°C for 24 hours under aerobic conditions in MRS broth 0.5% porcine bile (obtained from the Biological Services Unit (BSU), UCC) in a sterile 96-well plate. Negative controls contained MRS only and MRS with the corresponding bile of different origin but without bacteria. All samples were examined in triplicate and growth was monitored at 600nm over a 24-hour period by ThermoScientific Multiskan FC with SkanIt software 2.5.1. To assess survival isolates were passaged through predicted biological parameters encountered in the GIT, as described and reformulated by Vizoso Pinto *et al*., (2006)(Vizoso Pinto *et al*., 2006).

### Bile salt hydrolase activity assessment with UPLC-MS

Both wild type isolates and *bsh* clones were subjected to co-incubation assays in 0.5% porcine bile followed by UPLC-MS using methods described by Joyce *et al*.,(2014)(Joyce *et al*., 2014).

### Statistical analysis

Data analysis was carried out using Graphpad Prism version 5.0 software (https://www.graphpad.com/). Statistical tests were performed using one-way ANOVA followed by Dunnett’s post-hoc test unless stated otherwise. Results were considered statistically significant when p<0.05 (ns-no significance, One-way ANOVA: *= p<0.05; **=p<0.01; ***=p<0.001; ****=p<0.0001.). Where demonstrated, heatplots were generated using R version 3.6.0 and RStudio software version 1.1.453.

## Acknowledgements and Funding

SLL and SAJ designed the research and wrote the paper. SLL performed the research. We thank Mr Peter Cronin and Ms Christina Killian for technical assistance in the isolation of strains. This work was supported by Science Foundation Ireland Centres Grant SFI/12/RC/2273 P2 to APC Microbiome Ireland. S.A.J. and this work is also funded by SFI: EU Joint Programme Initiative CABALA for Health 16/ERA-HDHL/3358 and Ireland Department of Agriculture, Food and the Marine (DAFM) Award No. DAFM 17-RD-US-ROI.

